# Feasibility of constructing multi-dimensional genomic maps of juvenile idiopathic arthritis

**DOI:** 10.1101/349811

**Authors:** Lisha Zhu, Kaiyu Jiang, Laiping Wong, Michael J. Buck, Yanmin Chen, Halima Moncrieffe, Laura A. McIntosh, Kathleen M. O’Neil, Tao Liu, Xiaoyun Xing, Daofeng Li, Ting Wang, James N. Jarvis

## Abstract

**Background:** Juvenile idiopathic arthritis (JIA) is one of the most common chronic conditions of childhood. Like many common chronic human illnesses, JIA likely involves complex interactions between genes and the environment, mediated by the epigenome. Such interactions are best understood through multi-dimensional genomic maps that identify critical genetic and epigenetic components of the disease. However, constructing such maps in a cost-effective way is challenging, and this challenge is further complicated by the challenge of obtaining biospecimens from pediatric patients at time of disease diagnosis, prior to therapy, as well as the limited quantity of biospecimen that can be obtained from children,particularly those who are unwell. In this paper, we demonstrate the feasibility and utility of creating multi-dimensional genomic maps for JIA from limited sample numbers.

**Methods:** To accomplish our aims, we used an approach similar to that used in the ENCODE and Roadmap Epigenomics projects, which used only 2 replicates for each component of the genomic maps. We used genome-wide DNA methylation sequencing, whole genome sequencing on the Illumina 10x platform, RNA sequencing, and chromatin immunoprecipitation-sequencing for informative histone marks (H3K4me1 and H3K27ac) to construct a multi-dimensional map of JIA neutrophils, a cell we have shown to be important in the pathobiology of JIA.

**Results:** The epigenomes of JIA neutrophils display numerous differences from those from healthy children. DNA methylation changes, however, had only a weak effect on differential gene expression. In contrast, H3K4me1 and H3K27ac, commonly associated with enhancer functions, strongly correlated with gene expression. Furthermore, although unique/novel enhancer marks were associated with insertion-deletion events (indels) identified on whole genome sequencing, we saw no strong association between epigenetic changes and underlying genetic variation. The initiation of treatment in JIA is associated with a re-ordering of both DNA methylation and histone modifications, demonstrating the plasticity of the epigenome in this setting.

**Conclusions:** These findings, generated from a small number of patient samples, demonstrate how multidimensional genomic studies may yield new understandings of biology of JIA and provide insight into how therapy alters gene expression patterns.

## Introduction

Gene-environment interactions are thought to mediate many complex human traits, including human diseases [1], [2]. It is becoming increasingly clear that the influences of environment (broadly considered) are mediated through epigenetic changes to DNA and DNA-associated histones [3], [4], [5]. Thus, the field of epigenetics is emerging as an important complement to genetics for our understanding of complex traits [6], including rheumatic diseases such as rheumatoid arthritis [7] and systemic lupus erythematosus [8]. Juvenile idiopathic arthritis (JIA) is among those diseases that has been identified as a complex trait emerging from gene-environment interactions [9]. While substantial progress has been made in identifying genetic risk for JIA [10], our understanding of underlying epigenetic changes, and how/whether such changes contribute to disease risk or therapeutic response remains unknown. Furthermore, little is known, in the context of human diseases, about interactions between underlying genetic polymorphisms and the epigenome.

Our group has proposed that, rather than being understood primarily as an autoimmune disease, initiated by the faulty recognition of self-peptides by the adaptive immune system, JIA emerges because of complex interactions between innate and adaptive immunity [11]. We have demonstrated aberrant patterns of neutrophil activation that are linked to fundamental metabolic and oscillatory properties in JIA neutrophils [12] and reported aberrant patterns of gene expression associated with the metabolic abnormalities. [13],[14],[15] The aberrant patterns of gene expression do not return to normal when children are successfully treated [16]. Furthermore, contingency analyses of JIA neutrophil expression data [11] suggest a fundamental breakdown in those mechanisms that coordinate gene expression on a genome-wide basis [17].

In order to understand the interaction between the epigenome and underlying genetic variance, the field is going to require multi-dimensional genomic maps that query epigenetically-marked transcriptional regulators (e.g., H3K4me1/H3K27ac-marked enhancers), chromatin organizers (e.g., H3K9me3, CTCF binding sites), and other epigenetic modifiers of chromatin structure/accessibility (e.g., DNA methylation), and link those observations to observed aberrations in transcription. However, obtaining a sufficiently large volume of blood to perform such studies using single samples acquired from individual children is neither safe nor practical, and the expense of acquiring, processing, and analyzing a large group of samples adds another impediment.

We have therefore explored whether the necessary “genomic maps” might be generated using an approach similar to that used in the ENCODE and Roadmap Epigenomics projects: performing different analyses on small groups of cross-sectional samples and overlaying the data to provide a broad outline of functionally/ pathologically relevant regions. Such maps might then serve as a reference that allows other investigators to focus on specific genomic regions or chromatin features, which are not annotated in either ENCODE or Roadmap Epigenomics data sets, since those data sets carry no information about cells from human diseases or even healthy children. It should be noted that the Roadmap Epigenomics data sets, which have provided physician-scientists and geneticists a treasure trove of information that has been used to understand human disease, were generated using only two replicates [18]. We therefore set out to determine whether useful information might be derived from small numbers of replicates if multiple genomic modalities for pathologically relevant cells in JIA were overlayed in a manner similar to the Roadmap Epigenomics data.

To accomplish these aims, we started with neutrophils, a cell type that can be safely obtained in relative abundance from children and with which we have considerable experience. We performed genome-wide epigenome analyses using cross-sectional samples to build ENCODE-like epigenetic maps. We then queried whether such maps could be used to address relevant scientific and clinical questions, such as: (1) whether epigenetic risk for the disease is reflected in JIA neutrophils; (2) whether epigenetic alterations reflect underlying disease-associated genetic variance; and (3) whether epigenetic alterations are a permanent feature of JIA neutrophils; specifically, we asked whether treatment for JIA alters the neutrophil epigenome.

## Material and Methods

**Patients and patient samples for multi-dimensional analysis**. Neutrophils were extracted from all patients with the polyarticular, rheumatoid factor negative form of JIA as determined by internationally-accepted criteria [19]. (hereafter referred to as patients with JIA) and pediatric healthy controls.

**Whole genome methylation cohort:** This group consistented of 9 patients with JIA and five healthy control children (HC). Patients with JIA were grouped by disease state and initiation of methotrexate, the standard therapy for JIA. Specifically, we studied 6 patients with active, untreated disease (referred to as ADU) and 3 patients with active disease who had been started on methotrexate (,these patients are referred to as ADT). Disease activity was assessed using standardized criteria developed by Wallace and colleagues [20]. In addition, we performed whole genome methylation analysis on 5 healthy children (HC).

**ChIP-seq cohort:** Five children with JIA were studied: two children had untreated, newly-diagnosed JIA and 3 children were on therapy with methotrexate, studied 6 to 8 weeks after the initiation of therapy. We also studied 3 healthy controls.

**RNA-seq cohort:** Six children with JIA were studied and two healthy controls. Of the patients with JIA three children had untreated, newly-diagnosed JIA and three children had active JIA on therapy with methotrexate.

All patients with JIA were recruited from the pediatric rheumatology clinics at the University of Oklahoma, Cincinnati Children’s Medical Center, and the Women and Children’s Hospital of Buffalo. The healthy control (HC) children were recruited from the University of Oklahoma and Women and Children’s Hosopital of Buffalo general pediatrics clinics. All protocols for sample collection were reviewed and approved by the University of Oklahoma, University at Buffalo, and Cincinnati Children’s Hospital Institutional Review Boards. All research was carried out in compliance with the approved protocol. Informed consent documents were signed by the parents of all subjects, which also included consent to published de-identified information. Where appropriate, assent documents were signed by children over the age of 7 years in accordance with Cincinnati Children’s and University of Oklahoma IRB requirements.

**Cell isolation**. Neutrophils were separated by density-gradient centrifugation as described previously [12].In brief, whole blood was drawn into 10 mL CPT tubes (Becton Dickinson, Franklin Lakes, NJ), which is an evacuated blood collection tube system containing sodium citrate anticoagulant and blood separation media composed of a thixotropic polyester gel and a FICOLL™ Hypaque™ solution. Cell separation procedures were started within 1 h from the time the specimens were drawn. Neutrophils were separated by density-gradient centrifugation at 1,700× g for 20 min. After removing red cells from neutrophils by hypotonic lysis, neutrophils were then immediately treated differently depends on the downstream work. Cells prepared in this fashion are more than 98 % CD66b + by flow cytometry and contain no contaminating CD14+ cells, as previously reported [12]. Thus, although these cell preparations contained small numbers of other granulocytes, they will be referred to here as “neutrophils” for brevity and convenience. For RNA-seq, neutrophils were placed in TRIzol^®^ reagent (Invitrogen, Carlsbad, CA) and stored at −80 °C until used for RNA isolation. For DNA methylation sequencing study, neutrophil pellets were stored at −80 °C until used. For ChIP-seq study, neutrophils were immediately treated using 1% formaldehyde following the protocol (see description below) and then stored at −80 °C until used.

**MeDIP-seq and MRE-seq library generation, sequencing and mapping**. MeDIP and MRE sequencing libraries were generated as described previously [PMC4300244] with minor modifications. Briefly, for MeDIP-seq, 500 ng of DNA isolated was sonicated, end-processed, and ligated to paired-end adapters. After agarose gel size-selection, DNA was immunoprecipitated using a mouse monoclonal anti-methylcytidine antibody. Immunoprecipitated DNA was washed, amplified by 12 cycles of PCR, and size-selected by agarose gel electrophoresis. For MRE-seq, 5 parallel digests (HpaII, Hin6I, SsiI, BstUI, and HpyCH4IV; New England Biolabs) were performed, each with 200 ng of DNA. Digested DNA was combined, size-selected, end-processed, and ligated to single-end adapters. After the second size-selection, DNA was amplified by 15 cycles of PCR and size-selected by agarose gel electrophoresis.

MeDIP and MRE libraries were sequenced on Illumina HiSeq machine with a total number of approximately 1.28 billion MeDIP-seq reads and 583 million MRE-seq reads.

**RNA Isolation and Sequencing**. Total RNA was extracted using Trizol^®^ reagent according to manufacturer’s directions. RNA was further purified using RNeasy MiniElute Cleanup kit including a DNase digest according to the manufacturer’s instructions (QIAGEN, Valencia, CA). RNA was quantified spectrophotometrically (Nanodrop, Thermo Scientific, Wilmington, DE) and assessed for quality by capillary gel electrophoresis (Agilent 2100 Bioanalyzer; Agilent Technologies, Inc., Palo Alto, CA). Single-end cDNA libraries were prepared for each sample and sequenced using the Illumina TruSeq RNA Sample Preparation Kit by following the manufacture’s recommended procedures. And then sequenced using the Illumina HiSeq 2000. Library construction and RNA sequencing were performed in the Next generation sequencing center in University at Buffalo.

**Pre-processing and differential gene expression analysis of RNA-Seq data**. The raw reads obtained from paired-end RNA-Seq were mapped to human genome hg19, downloaded from the University of California Santa Cruz Genome Bioinformatics Site (http://genome.ucsc.edu) with no more than 2 mismatches using tophat v2.0.10 [21]. Gene expression level was calculated as FPKM (Fragments Per Kb per Million reads) with Cufflinks v2.1.1 [22], the annotation gtf file provided for genes is gencode v19 annotation gtf file from GENCODE (http://www.gencodegenes.org) [23]. We used Cuffdiff v2.2.1 [24] for pairwise comparisons of ADU, ADT and HC to identify differentially expressed genes (DEGs), with a false discovery rate of 5%. Genes with average FPKM >=1 in at least one of the three groups were considered as expressed genes in neutrophils.

**Chromatin immunoprecipitation for histone marks H3K4me1 and H3K27ac and sequencing**. Neutrophils were isolated as described previously [16]. The ChIP assay was carried out according to the protocol of manufacturer (Cell Signaling Technologies Inc., Danvers, MA, USA). Briefly, adult neutrophils were incubated with newly prepared 1% formaldehyde in 10 ml PBS for 10 min at room temperature (RT). Crosslinking was quenched by adding 1× glycine and incubation for 5 min at RT. The crosslinked samples were centrifuged at 800 x g for 5 min. The supernatant was discarded, and the pellet was washed two times with cold PBS followed by resuspension in 10 ml ice-cold Buffer A plus DTT, PMSF and protease inhibitor cocktail. Cells were incubated on ice for 10 minutes and centrifuged at 800 x g for 5 min at 4°C to precipitate nuclei pellets, which were then resuspended in 10 ml ice-cold Buffer A plus DTT. The nuclei pellet was incubated with Micrococcal nuclease for 20 minutes at 37°C with frequent mixing to digest DNA to lengths of 150 – 900bp. Sonication of nuclear lysates was performed using Sonic Dismembrator (FB-705, Fisher Scientific, Pittsburgh, PA, USA) on ice under the following conditions: power of 5, sonication time of 30 seconds with pulse on 10 s and pulse off 30 s. After centrifugation of sonicated lysates at 10000 x g at 4°C for 10 min, the supernatant was transferred into a fresh tube. Fifty microliters of the supernatant (chromatin preparation) was taken to analyze chromatin digestion and concentration. Fifteen micrograms of chromatin was added into 1 x ChIP buffer plus protease inhibitor cocktail in a total volume of 500 μl. After removal 2% of chromatin as input sample, the antibodies were added in the ChIP buffer. The antibodies against respective histone modifications are: rabbit polyclonal antibodies against histone H3 acetylated at lysine 27 (H3K27ac) and histone H3 monomethylated at lysine 4 (H3K4me1) from Cell Signaling Technologies (Danvers, MA, USA). The negative control is normal IgG (Cell Signaling Technologies. Danvers, MA, USA). After immunoprecipitation (IP) overnight at 4°C, the magnetic beads were added and incubated another 2 hours at 4°C. The magnetic beads were collected with magnetic separator (Life Technologies, Grand Island, NY, USA). The beads were washed sequentially with low and high salt wash buffer, followed by incubation with elution buffer containing proteinase K and NaCl to elute protein/DNA complexes and reverse crosslinks of protein/DNA complexes to release DNA. The DNA fragments were purified by spin columns and dissolved in the elution buffer of a total volume of 50 μl. The crosslinks of input sample were also reversed in elution buffer containing proteinase K before purification with spin columns. Then DNA-sequencing was conducted using the Illumina HiSeq 2500 at the next generation sequencing center in University at Buffalo.

**ChIP-Seq analysis of neutrophils**. Sequencing reads from ChIP-Seq experiments were mapped to human genome hg19 using BWA (Burrows-Wheeler Aligner, version 0.7.7-r441) [25]. MACS2 v2.1.10 [26] was applied for calling regions enriched with histone marks against the input sample, with the parameters “–nomodel --extsize 150 --broad -broad-cutoff 0.1”.

**Genomic features**. The broad regions of H3K4me1 and H3K27ac were annotated using CEAS software [27]. The annotation of a given region is decided by asking which of the following genomic features in order can be first overlapped: the 3 kbps upstream of known transcription start site as “promoter”, the 3kbps downstream of transcription termination site as “downstream”, “3’ UTR”, “intron”, ‘5’ UTR”, “exon” or the “intergenic” regions.

**Identification of differentially enriched regions (DERs)**. To identify differentially enriched regions, H3K27ac and H3K4me1 ChIP-Seq data were grouped according to disease conditions: ADT (n=3), ADU (n=2) and HC (n=3). The peak regions obtained from MACS2 v2.1.10 [26] in each individual of three groups were combined together, sequence coverage of each individual was calculated using MACS2 v2.1.10 [26] pileup by extending each read to length of 150 bp and the resulting coverage profiles were exported as bigwig files. The peak regions were binned to 200-bp windows with a sliding window of 100 bp, the enrichment values which expressed as reads per kilobase per million mapped (RPKM) were calculated for each 200-bp windows by using bigwig files. edgeR [28] was then applied to discover initial DERs between ADU and HC, the P-value and fold-change (FC) threshold were decided according to volcano plot. (FDR < 0.001 for H3K27ac and FDR < 0.05 for H3K4me1 in ADU vs. HC; FC cutoff was set to 2). The DERs exhibited distinct variability between ADU and HC for both H3K27ac and H3K4me1, demonstrating distinct clusters that allowed accurately classification of each sample (Figure 1a). MACS2 bdgbroadcall [26] was applied to obtain the broad DERs which were used in further analysis from the initial DERs. The lower cutoff of FDR was set to 0.05 for H3K27ac and 0.1 for H3K4me1, the lower cutoff of FC was set to 1.5. The initial DERs between ADU and ADT were obtained using a P value < 0.005 and FC > 2 for both H3K27ac and H3K4me1. To call broad DERs, the lower cutoff of P value was set to 0.05 and FC was set to 1.5. The intersection of broad DERs gained or lost in the ADU group in both ADU vs. HC and ADU vs. ADT were considered as regions significantly affected by treatment. To obtain the treatment-unique DERs, we treated ADU and HC together as one group and compared them with ADT. The initial DERs were obtained from edgeR [28] using P value < 0.001 and FC > 2. For broad DER calling, the lower cutoff for P value was 0.05 and 1.5 for FC. The broad DERs obtained by the method were then intersected with the regions considered as similar between ADU and HC (log2FC < 0.1 as peak cutoff and log2FC < 0.2 as linking peaks cutoff in MACS2 bdgbroadcall) to obtain ADT unique DERs. All the DERs used for further analysis should with length no less than 200bp.

**Figure 1.**
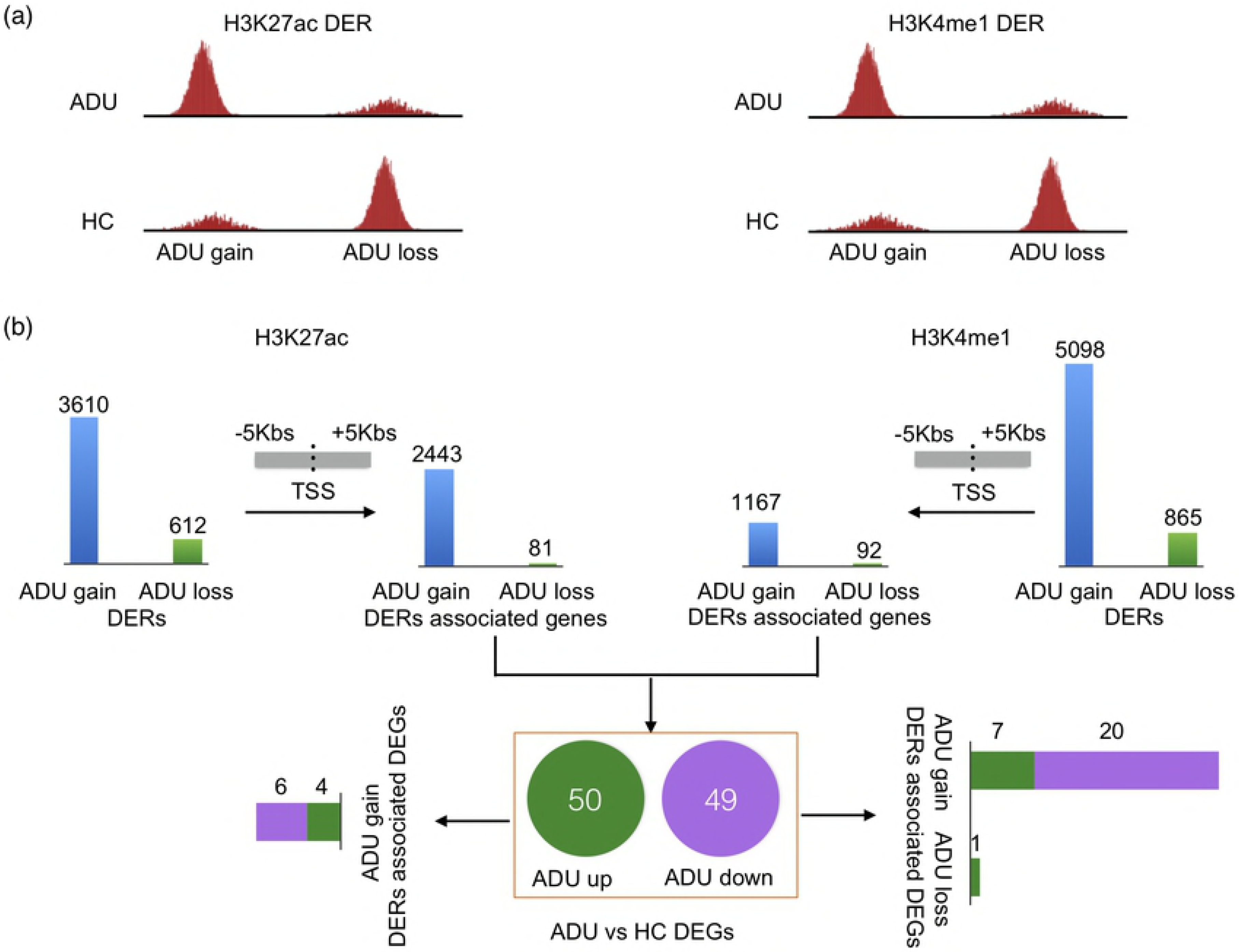
Graphic illustration of definitions and processing steps to identify DERs between ADU and HC. (a) Graphic sketch to show the definition of ADU H3K27ac/H3K4me1 gain and ADU loss DER, ADU gain DER has much higher histone mark signals than ADU loss DER. (b) A flowchart to show the steps to associate DERs with DEGs. Those genes with upstream 5 Kbs to downstream 5 Kbs of TSS contained DERs are defined as DERs associated genes, those genes further compared with DEGs between ADU and HC to obtain DERs associated DEGs.

**Identification of differentially methylated regions (DMRs)**. The reads of MeDIP-Seq and MRE-Seq were aligned with bwa [25] to human hg19. Female and male samples were mapped with female and male bwa index separately. Regions related to X and Y chromosomes were removed to avoid sex bias. Pairwise statistical analyses of DMRs in ADU (n=6), ADT (n=3) and HC (n=5) groups were performed with repMnM, a modified version of M&M [29]. The bin length was set to 500bp, and those regions with q values smaller than 1×10^−2^ were selected as significant DMRs.

**Whole Genome Sequencing (WGS)**. We performed WGS on the Illumina X Ten platform on 50 samples from 48 individuals with JIA (duplicates were sequenced to assess quality and fidelity of the sequencing data). Of these, 37 subjects were girls and 11 were boys. All patients were of European descent (n=33) or mixed European-American Indian descent (n=5). We have recently published a preliminary analysis of these data [30].

We used BWA [25] to map sequencing reads of each sample with respect to the human reference genome hg19. Potential artifacts caused by PCR duplicated reads were removed using Picard tools (http://picard.sourceforge.net.). Genome Analysis Toolkit (GATK version 3.2-2-gec30cee) ^36^ was used to refine reads around small insertions or deletions (indels) and recalibrate base quality scores. We adopted the GATK best practices in calling SNPs, small insertion or deletion (indels) between 1 and 50 bp in size. The first step of SNP and indel discovery involved single sample variant calling (GATK HaplotypeCaller), followed by joint genotyping per cohort (GATK GenotypeGVCFs) and variant quality variant score recalibration (VQSR, GATK VariantRecalibrator). In order to discover high quality variants, we retained variants that passed variant filtration criteria consisting of minimum read depth of 20X, genotype quality ≥20, variant quality ≥30 and minor-read ratio (MRR) ≥0.2. MMR was accessed to ensure the read of the less covered allele (reference or alternate) over the total number of reads have sufficient depth to be called as a variant. The variant filtration criteria aim to minimize false positives in genetic variant discovery. We note that in the 48 WGS on JIA samples, we performed two batches of sequencing (29 samples in the first batch and 19 samples in the second batch) to validate variant discovery. From the variant concordance analysis performed on the 2 sequencing batches, we found a concordance rate >83% for SNPs and indels, indicating that the detected genetic variants are likely to be authentic and suitable for downstream analysis.

**Interrogating with whole genome sequencing DNA variants of 48 JIA individuals**. In order to discover the possible regulation relationship between enhancers and genomic variation, we intersected the detected DER (DMR) with variants discovered from deep whole genome sequencing (WGS) of 48 JIA individuals using bedtools program[8]. Next, we adopted Fisher exact test for enrichment analysis to inform the statistical significance for the association between DER (DMR) and genetic variants relative to genetic variants of the 1000 Genomes Projects (1KGP). We used this approach as it would be impossibly expensive to conduct WGS on a healthy control cohort. Furthermore, the whole purpose of publicly available healthy genome sequences is to serve for a basis of comparisons in studies like this one. At the sample collection time, 2504 subjects from 1KGP declared themselves to be healthy. We therefore tested the difference in proportion of SNPs overlapping DER (DMR) between JIA SNPs and 1KGP SNPs. First, we constructed 2 by 2 contingency tables, with the first row containing the number of overlapping SNPs (with DER or DMR) for JIA (first column) and 1KGP (second column), while the second row contained numbers of non-overlapping SNPs for JIA and 1KGP. Fisher exact test’s p-value was computed to gauge the significance of the observed differences. The odds ratio of the Fisher exact test was used as enrichment fold. Using this approach, we tested whether there were more genetic variants derived from JIA genomes compared to genetic variants in the 1KGP data set when overlapped with epigenetic elements (DER or DMR). All analyses were conducted using in-house customized Perl and R scripting under Linux environment.

## Public Availability of Genomic Data

We have made the data available to the scientific community. WGS bam files have been uploaded to NCBI bioproject: https://www.ncbi.nlm.nih.gov/bioproject/?term=PRJNA343545.

WGS snps/indels were uploaded to dbsnp149: handle id: JJLAB. WGS structural variatiants were submitted to DGVa id: estd231 https://www.ncbi.nlm.nih.gov/dbvar/studies/estd231/.

Other data were uploaded to the Gene Expression Omnibus (GEO). The GEO accession number for the RNA-Seq in neutrophils is GSE92293, for ChIP-Seq of H3K27ac and H3K4me1 in neutrophils is GSE92393, and for DNA methylation in neutrophils is GSE92749.

## Results

As noted in our Introduction, our goal in this study was to test the feasibility of generating high-quality maps of the epigenomes of pathologically relevant cells in disease states, mirroring the Roadmap Epigenomics and ENCODE data. Such maps might then be used to guide focused studies on specific regions of interest, for example, in studies aimed to understand the relationship between genetic variation and chromatin structure, or studies aimed at understanding mechanisms underlying the transcriptional abnormalities observed in peripheral blood cells of children with JIA [15, 31].

### Differentially expressed genes from RNA-Seq analysis

We first sought to corroborate our previous observations that JIA neutrophils show distinct transcriptional abnormalities compared to neutrophils in healthy children, and that treatment is associated with transcriptional reorganization [12, 14]. We identified 99 DEGs (FDR < 0.05, Fold change < 1.5) between ADU (3 individuals) and HC groups (2 individuals), with 50 genes showing higher expression in ADU and 49 genes showing higher expression in HC. Using all genes expressed in neutrophils as background, gene ontology (GO) analyses by the GOrilla tool [32] predictably showed these DEGs were enriched for immune effector processes and defense responses. To investigate whether the initiation of therapy with methotrexate was associated with transcriptional changes in patients with JIA, we also compared the gene expression between untreated (ADU) patients and children who had started methotrexate therapy 2-6 weeks earlier but still had active disease (ADT; 3 individuals). We identified 703 DEGs that demonstrated significant expression changes between ADU and ADT, with 341 genes showing higher expression levels in ADU and 362 showing higher expression in ADT. This extensive alteration in neutrophil transcription after the initiation of therapy corroborates our findings from hybridization-based microarrays [33]. GO analysis showed that the highly expressed genes in ADU were associated with cellular protein modification processes and regulation of cellular biosynthetic processes, and those with higher expression in ADT were enriched in protein targeting to the endoplasmic reticulum and viral transcription and translation. Lists of DEGs and enriched GO terms for each group are provided in **S.Table1**. These findings are consistent with what we have previously observed in JIA neutrophils using hybridization-based-microarrays, (i.e., that JIA neutrophils show aberrant patterns of transcription and that therapy re-orders, but does not normalize, neutrophil transcriptomes). [12]

We next investigated how expression levels of the 99 specific DEGs identified in the ADU vs. HC comparison might be altered during treatment. We found that, among the 99 DEGs identified in the ADU vs. HC comparison, 79 changed after initiation of treatment toward levels seen in HC, i.e., these genes have the same up or down fold-change trends in the ADU vs. ADT comparison (Figure S1). Thus, while the initiation of therapy begins to “correct” genes that show aberrant patterns of expression in untreated JIA, the ADT state has its own distinct transcriptional signature that differs both HC and ADU. These findings corroborate what we have previously reported from hybridization-based microarray studies [13].

In addition to corroborating our previous findings, the RNA-Seq studies show that reference “maps” of disease-specific epigenomes will need to be generated for specific disease states (e.g., untreated disease, disease that is active, disease that is in remission, etc). These studies further highlight the desirability of creating these maps from small numbers of samples in order to reduce the expense of generating them.

### Presence of disease-specific enhancers within JIA neutrophils

Our next step was to examine specific epigenetic marks to determine whether epigenetic signatures generated from patients might differ from reference sets. This is important, as current assessments of the link between genetic risk and epigenetic signatures, for example, have relied entirely on the analysis of data generated from healthy adults [34, 35].

We studied neutrophils collected from children with active, untreated JIA (ADU: 2 individuals) and healthy controls (HC: 3 individuals). We first used ChIP-Seq to study two histone marks, H3K27ac and H3K4me1, typically associated with enhancer activity. We binned the human genome, then used edgeR [28] to call H3K4me1 and H3K27ac DERs with significant and robust alterations in ChIP-Seq signals (See Methods for detail). The definition and analysis of DERs between ADU and HC are shown in Figure 1a, 1b. We identified 3,610 (average length: 3610 bps) H3K27ac and 5,098 (average length: 781 bps) H3K4me1 DERs gained in the ADU group (i.e., regions that were more enriched with histone modifications in the ADU than in the HC group). We also identified 612 (average length: 1006 bps) H3K27ac and 865 (average length: 865 bps) H3K4me1 DERs lost in the ADU group (Figures 2a, 2b) (i.e., regions that were more enriched for H3K4me1 or H3K27ac in the HC group than in the ADU group). We note that ADU gained both more H3K27ac DERs and H3K4me1 DERs compared to HCs. The ADU loss DERs are significantly enriched in introns and depleted in intergenic regions when compared to genomic background; while the ADU gained DERs are significantly enriched in promoter regions (Figure 2c). Specifically, more than 90% of the H3K4me1 loss DERs in ADU are located within introns.

**Figure 2.**
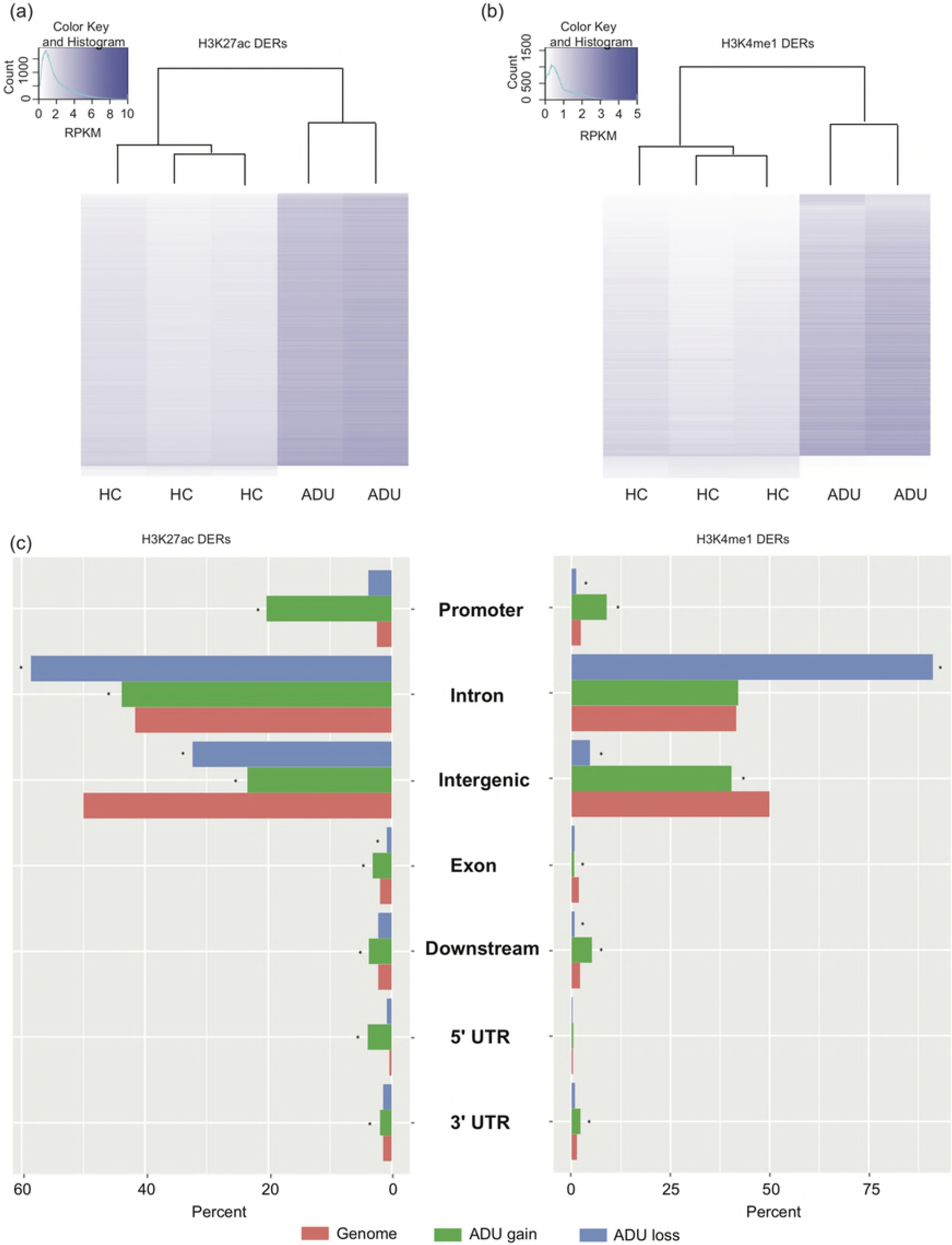
Heatmap and genomic distribution of H3K27ac and H3K4me1 DERs between ADU and HC. (a) Heatmap of H3K27ac RPKM values. (b) Heatmap of H3K4me1 RPKM values. (c) Genomic distribution of H3K27ac and H3K4me1 DERs between ADU and HC. (Fisher’s exact test for the difference between DERs and genome background, * P < 10^−6^).

We next investigated the DERs within gene promoter regions and non-promoter regions separately (we considered the regions from 5Kbs upstream to 1Kbs downstream around the transcription start site (TSS) to be promoter regions and designated other regions as non-promoter regions). The gained DERs were classified into three categories: (1) regions having both H3K27ac and H3K4me1 gains; (2) regions having only H3K27ac gains; or (3) regions having only H3K4me1 gains. Regions gaining both H3K27ac and H3K4me1 marks typically indicate increasing enhancer activity [36], so we initially focused on these regions. We found that, in both promoter and non-promoter regions, ADU gained more and lost fewer such DERs with both H3K27ac and H3K4me1 marks (**S. Table2**). Taken together, these findings corroborate other studies that disease states are accompanied by significant alterations in the functional epigenomes of disease-relevant cells and highlight again the desirability of developing disease-specific “reference epigenomes” to complement data generated from healthy individuals.

### Enhancer aberrations correlate with transcriptional alterations in JIA neutrophils

We next examined the regulation potential of enhancer changes in JIA neutrophils using BETA software. [37] The regulation potential for a certain gene is a summary of both the number of regulatory elements nearby and the distances (±5Kbs used in the study) to TSS. ADU gain H3K27ac DERs and H3K4me1 DERs both demonstrated activating and repressive functions on gene expression (p value < 10^−6^). While ADU loss H3K27ac and H3K4me1 DERs showed no significant regulatory functions, this might be related to the small number of DERs in the two groups. We then identified DER-associated genes, also using windows of −5Kbs to +5Kbs around the TSS, and intersected these regions with our detected DERs, only those genes considered to be expressed in neutrophils were kept (see Methods and (**S.Table3**)). Using GO term analysis with GOrilla software [32] and the genes expressed in neutrophils as background, we found that the genes associated with ADU H3K27ac gained DERs were enriched for regulation of metabolic and biological processes. This finding is of interest given our previous report that JIA neutrophils display disordered regulation of metabolically-mediated oscillatory functions that are critical to myeloperoxidase and superoxide ion production [13]. The genes associated with H3K4me1 DERs gained in ADU were enriched for GO terms for the regulation of immune system processes, cell migration and cell adhesion. These processes are fundamental to neutrophil phagocytic functions, and we have recently identified aberrant patterns of expression for genes associated with cell migration and adhesion in whole blood expression data from children with untreated JIA [31]. Genes that associated with H3K27ac DERs loss in ADU were significantly enriched for immune system processes, regulation of cell activation and cell adhesion (**S.Table4**). All the DEGs which are associated with DERs are shown in **S.Table5**. Taken together, these findings demonstrate that alterations in H3K4me1/H3K27ac histone marks are associated with differential transcription in JIA neutrophils, and the identified H3K27ac and H3K4me1 alterations have observable effects on gene transcription as identified on RNA-Seq.

In contrast, the DMRs identified between ADU and HC were not significantly associated with differential gene expression when considering hyper- and hypo-methylated regions separately. We designated the genes that were (−5Kbs, 5Kbs) of TSS or whose gene bodies intersected with DMRs as DMR-associated genes. We identified 160 expressed genes associated with hypermethylated DMRs and 100 expressed genes associated with hypomethylated DMRs in the ADU group (**S.Table6**). However, GO analyses did not reveal any significantly enriched terms in biological processes. Only three ADU hypermethylated DMR-associated genes (*ADARB2*, *CNTNAP3B*, and *MYOM2*) showed significant differential expression in the ADU vs. HC comparison of RNA-Seq data. It should be noted that some of the DMRs are located within the H3K27ac and H3K4me1 enhancer regions (**S. Table7**) and may interact with enhancer elements and thus impact gene transcription indirectly.

### Therapy is associated with alterations in H3K4me1/H3K27ac-defined enhancer marks within JIA neutrophils

We next sought to determine whether epigenetic features seen in JIA neutrophils are a permanent feature of these cells. To accomplish this aim, we examined whether treatment with methotrexate, a first-line therapy for polyarticular JIA, impacts the epigenetic signatures observed in JIA neutrophils. For all H3K27ac or H3K4me1 DERs that we identified in the comparison of the ADU with HC groups, we examined the H3K27ac and H3K4me1 signal intensities within those DERs in children who had been on standard therapy with methotrexate for 6-8 weeks but still had active disease (designated the ADT group). The majority of ADU gain H3K27ac or H3K4me1 DERs (compared to HC) showed decreased H3K27ac or H3K4me1 marks after treatment (Figure S3a,3b). We identified 470 (average length: 705 bps) DERs gained H3K27ac in ADU compared with HC but then lost H3K27ac in ADT, and 818 (average length: 460 bps) gained H3K4me1 in ADU then lost in ADT (Figure 3a). In other words, after treatment, these regions gaining the histone marks in the active disease state changed significantly toward levels identified in HC (Figure 3c, 3d); we designate these as “gain-then-loss” DERs. On the other hand, a smaller number of DERs (33 for H3K27ac and 13 for H3K4me1) gained H3K27ac or H3K4me1 in the ADU compared with HC and showed a further, significant H3K27ac or H3K4me1 gain in ADT vs ADU. We also detected the “loss-then-regain” DERs, that is, 55 (average length: 494 bps) H3K27ac DERs lost in ADU but then regain the mark in ADT and 140 (average length: 322 bps) DERs lost H3K4me1 in ADU then regain it in ADT (Figure 3b). These regions also demonstrated histone marks that were more similar, compared with the gain-then-loss DERs, to HC after the initiation of treatment (Figure 3c, 3d). Only a minority (4 for H3K27ac and 5 for H3K4me1) of DERs that lost H3K27ac or H3K4me1 showed a significant loss again in ADT. Taken together, these results demonstrate that after treatment is initiated, both H3K27ac and H3K4me1 marks are changed and more closely resemble the pattern seen in HC, especially for those enhancers that lost activity in active disease. The genes associated with those DERs were displayed in **S. Table8** and show similar expression levels in ADU, HC and ADT groups (Figure S3c,3d).

**Figure 3.**
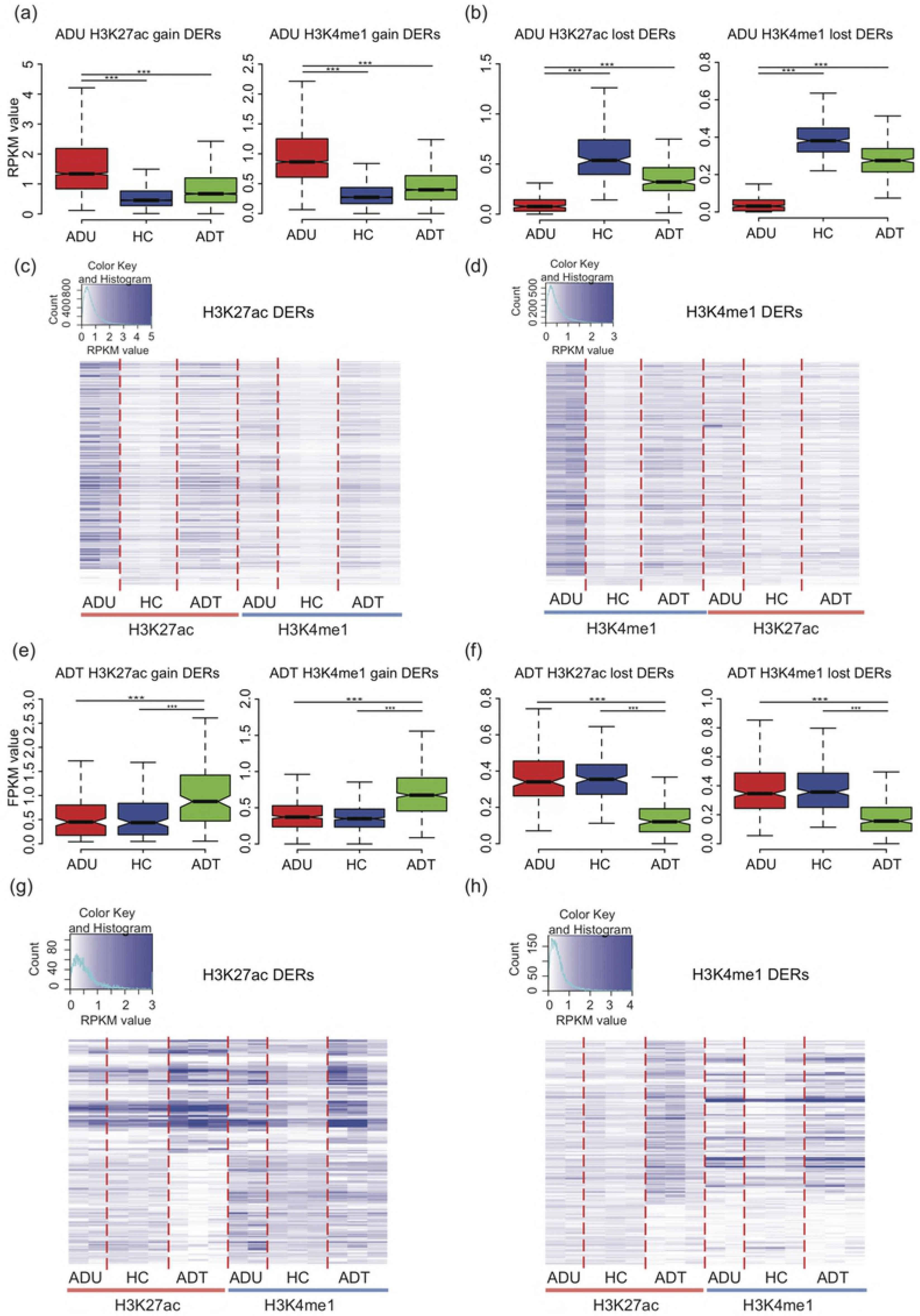
Treatment with methotrexate alters JIA neutrophil epigenomes. H3K27ac and H3K4me1 marks among ADU, HC and ADT of DERs identified in the comparison between ADU and HC which also changed significantly between ADU and ADT. Treatment also exhibits unique H3K27ac and H3K4me1 marks of DERs identified as significantly different between ADT and ADU & HC. (a) boxplot of H3K27ac levels of ADU H3K27ac gain DERs (left) and H3K4me1 levels ADU H3K4me1 gain DERs (right) (Fisher’s exact test between ADU and HC, ADU and ADT; ‘*’ indicates 10^−4^ < P ≤ 10^−2^; ‘**’ 10^−15^ < P ≤ 10^−4^; ‘***’ P ≤ 10^−15^). (b) boxplot of H3K27ac levels of ADU H3K27ac loss DERs (left) and H3K4me1 levels ADU H3K4me1 loss DERs (right). (c) Heatmap of H3K27ac and H3K4me1 marks of 268 H3K27ac DERs between ADU and HC which were also significantly changed between ADU and ADT. (d) Heatmap of H3K4me1 and H3K27ac marks of 778 H3K4me1 DERs between ADU and HC which were also significantly changed between ADU and ADT. (e) boxplot of H3K27ac levels of ADT H3K27ac gain DERs (left) and H3K4me1 levels ADT H3K4me1 gain DERs (right). (f) boxplot of H3K27ac levels of ADT H3K27ac loss DERs (left) and H3K4me1 levels ADT H3K4me1 loss DERs (right). (g) Heatmap of H3K27ac marks of 488 ADT unique H3K27ac DERs. (h) Heatmap of H3K4me1 marks of 1509 ADT unique H3K4me1 DERs.

### Therapy exhibits unique H3K4me1/H3K27ac-defined enhancer marks within JIA neutrophils

From the RNA expression analysis, we have shown that the disease related genes (DEGs between ADU and HC) change their expression with treatment, although there are more genes differentially expressed in ADT compared with ADU. We hypothesized that therapy has a direct impact on patients’ neutrophils, possibly related to therapy-induced alterations in the epigenome. . In order to determine the treatment unique DERs (unique epigenetic changes produced by therapy), first we obtained the regions of those genomic loci having consistent histone marks (with log_2_(FC) < 0.1) between ADU and HC, then we obtained the DERs within these regions showing changed histone marks in ADT compared to non-ADT groups (both ADU and HC). We found that ADT gained 38 (average length: 521 bps) H3K27ac DERs and 136 (average length: 472 bps) H3K4me1 DERs (Figure 3e), and lost 47 (average length: 440 bps) H3K27ac DERs and 93 (average length: 417 bps) H3K4me1 DERs compared with the non-ADT groups (Figure 3f). This finding indicates that, after treatment, the histone marks of these regions changed to a unique state other than ADU and HC (Figure 3g, 3h). The expressed genes associated with those ADT unique DERs are displayed in **S.Table 9**.

Taken together, these findings demonstrate that, even for functional signatures such as H3K4me1/H3K27ac, JIA neutrophil epigenomes are quite plastic and reordered by therapy. They point to the promise that building multi-dimensional genomic maps across a range of relevant cells may elucidate vexing questions surrounding treatment response and refractoriness in JIA.

### Genome-wide DNA methylation in JIA neutrophils: presence of disease-associated methylation marks

Both ENCODE and Roadmap Epigenomics project annotate DNA methylation across a broad spectrum of cell types. DNA methylation is an important epigenetic regulator of gene expression, and it is interesting to note that the first-line therapy for JIA, methotrexate, is thought to inhibit methyltransferase activity. Thus we compared genome-wide DNA methylation using MeDIP-Seq and MRE-Seq assays, in neutrophils of ADU patients, with genome-wide methylation patterns seen in HCs. As with histone-associated epigenetic marks, we identified distinct disease and disease-state methylation signatures in JIA neutrophils.

We identified 1,072 differentially methylated regions (DMRs), of 500bps each, when comparing ADU with HC subjects. Of these, 705 were hyper-methylated and 367 were hypo-methylated in the ADU group (Figure S2a). These DMRs were enriched in distal intergenic regions and depleted in introns when compared to genomic background (Figure S2b, c), showing an opposite distribution compared to the two histone marks, H3K4me1 and H3K27ac. The methylation analysis showed that methylation changes between disease and healthy samples, supporting the idea that disease-specific data is likely to be useful in attempts to understand links between genetic variation and the epigenome. We next explored that question further.

### Link Between Genetics and Epigenetics

We next sought to determine the degree to which underlying genetic variation might determine the epigenetic signatures we observed in JIA neutrophils. We started by examining regions of known genetic risk. We queried whether there was enrichment for enhancer mark changes in linkage disequilibrium (LD) blocks that encompassed the 40 JIA-associated SNPs from the non-coding genome identified by Hinks et al [10]. We obtained the LD blocks of those SNPs from the SNAP database (http://www.broadinstitute.org/mpg/snap) reported by the 1000 Genomes Project pilot1 and HapMap3 projects with the cutoff of r^2^ <0.9. We then identified 26 LD blocks containing 35 out of the 40 SNPs identified from the genetic fine mapping study. Some SNPs were within the same LD block, and 5 SNPs cannot be associated with any LD blocks. We found that within LD blocks containing JIA-associated SNPs, compared with HC, only a limited number of ADU gained or lost DERs were found. Among them, the 5 LD blocks with ADU gained H3K27ac DERs and 6 LD blocks with ADU gained H3K4me1 DERs show statistical significance with p value < 0.01 (permutation test by randomly selecting 26 other genomic regions with the same length of the 26 LD blocks 10000 times and the average number of those 26 regions with ADU gained H3K27ac DERs is significant smaller than 5 and with ADU gained H3K4me1 DERs is significant smaller than 6). (**S.Table10**). We also investigated whether any of the differentially methylated regions (DMRs) identified in JIA neutrophils were located within the 26 LD blocks harboring JIA-associated SNPs, but found none.

These findings point to the utility of this approach. The regions where known genetic risk for JIA is associated with alterations in the functional epigenome will be of special interest to investigators interested in the mechanisms through which genetic variance alters non-coding genome function. Furthermore, these data provide a map that will sharply reduce the proportion of the involved haplotypes that need to be investigated (and therefore the number of SNPs that need to be tested) in functional assays. They also suggest that, at least in neutrophils, DNA methylation may not be an informative area of inquiry for studies aimed at understanding genetic effects on epigenetic alterations in JIA.

### Overlapping of regions with differentially enriched histone marks and DNA methylation with whole genome DNA sequencing (WGS) variants

We next sought to determine whether underlying epigenetic differences observed in the neutrophils of children with JIA (when compared to HC) might reflect underlying genetic variation beyond the established JIA haplotypes. Although GWAS and genetic fine mapping studies have identified regions of genetic risk for JIA [10], such studies have not truly been “genome-wide.” For example, the Hinks study [10] used the Illumina Immunochip, which queries 195,806 SNPs and 718 small insertions-deletions in regions of the genome that are of specific immunologic interest. In order to gain a broader understanding of genomic regions that exert effects on transcriptional control (including *trans* effects), we performed WGS on 48 children with JIA at an average sequencing depth of 39x (range 32-49x) using the Illumina X Ten platform at the New York Genome Center.

We took all categories of DER from either H3K27ac or H3K4me1 marks identified from the comparison between ADU and HC, considered their locations at promoter and non-promoter sites, and intersected these regions with JIA WGS variants (10,800,221 SNPs and 1,177,966 indels). We found that 24.72% and 23.73% of H3K27ac/H3K4me1 gain enhancer signals at promoter and non-promoter regions respectively co-localized with indels (**S.Table11**). Also, we found an average of 78.97% DERs (number of overlapping DERs range between 7 and 344, **S.Table11**) across different categories at promoter sites co-localized with SNPs. For 6 categories of DER at non-promoter sites, we identified between 2 to 694 overlapping DERs, with an average of 76.29% DERs co-segregating with SNPs (all SNPs include known and novel SNPs, **S.Table11**). By using Fisher’s Test with a cutoff p-value of ≤0.01, enrichment fold ≥1.5, and genetic variants of the 1000 Genomes Projects as background comparison, we did not observe enrichment of JIA variants within any of the 6 categories (**S.Table11**) of the reported DERs.

Next, we interrogated the DERs that showed potential changes after treatment by examining the H3K4me1/H3K27ac signal intensity for DERs (ADU vs HC) in children who still had active disease (ADT). This yielded an average of 18.49% DERs with potential changes due to therapy which overlapped with indels from WGS in JIA patients. In total we found between 9 and 164 DERs from 4 categories overlapping JIA indels (**S.Table12**). On the other hand, we observed that an average of 72.29% DERs with potential changes due to therapy overlapped with SNPs from WGS on JIA patients. As tabulated in **S.Table12**, we detected 44 to 568 DER across 4 categories overlapping with SNPs. We did not see any JIA genetic variant enrichment (or depletion relative to genetic variants of the 1000 Genomes Projects) in those DERs where we observed dynamic changes associated with treatment (**S.Table12**). Thus, underlying genetic variance was not strongly associated with differences in histone marks (at baseline or after therapy) in JIA neutrophils.

### Overlap of differentially DNA methylated regions between patients and controls with whole genome DNA sequencing (WGS) variants

We further determined whether differential DNA methylation as seen in JIA neutrophils could be accounted for by genetic variance between children with JIA and the populations represented the 1,000 Genomes Project. This analysis yielded 11% to 23% for different categories of DMR in ADU co-localized with indels found from WGS of 48 JIA patients (**S.Table13**). Also, we found 77%-82% of DMRs co-segregated with SNPs discovered from WGS of JIA individuals. Enrichment analysis using the Fisher exact test on each category of the DMR (hyper or hypo methylated in ADU at non-promoter or promoter regions) was conducted to determine whether any DMR category is enriched with JIA genetic variants. We found no evidence of enrichment with Fisher exact test p-value ≤0.01 and enrichment fold ≥1.5 (**S.Table13**). Overall, our findings fail to support the idea that underlying genetic variance accounts in any significant way for the observed epigenetic differences in JIA neutrophils.

Taken together, these small, independently-acquired data sets do not indicate a strong association between genetic variation (as identified by the WGS data) and epigenetic alterations in JIA neutrophils. It seems likely that specific, directed experiments will be required to identify allelic effects on epigenetic signals at specific genomic locations[38].

## Discussion

In the current work, we demonstrate the feasibility of developing informative, multidimensional genomic maps of JIA to study the interplay between genetics and epigenetics in this common childhood disease. We took the same approach used by the NIH ENCODE and Roadmap Epigenomics projects, seeking to gather the largest amount of information from the smallest number of samples as possible. We started by assuring ourselves that this parsimonious approach could recapitulate findings we have already published: i.e., that neutrophil transcriptomes are abnormal in JIA neutrophils and change over the course of therapy [12-16]. Having corroborated our previous work using a small number of samples for RNA-Seq, we proceeded to explore the kinds of information that might emerge from building a multi-dimensional genomic model from cross-sectional samples.

Our current work corroborates previous investigators’ findings that there are disease-specific epigenetic changes that can be observed in peripheral blood cells of patients with chronic inflammatory diseases. For example, Jeffries et al reported distinct patterns of DNA methylation in CD4+ T cells of patients with systemic lupus erythematosus, which included hypomethylation of genes known to be involved in T cell activation and implicated in autoimmunity [39]. More recently, Seumois et al reported the presence of novel H3K4me1/H3K27ac-marked enhancers in primary human T cells. These enhancers were associated with asthma susceptibility and correlated with asthma-associated SNPs [40]. Our work here corroborates the idea that there are disease-specific alterations in the functional epigenome as indicated by H3K4me1/H3K27ac (enhancer) marks that were seen only in JIA neutrophils, as well as the absence of such marks in JIA neutrophils in regions where they are seen in healthy controls. The fact that we can corroborate findings from studies that were generated using larger numbers of samples supports the utility of generating such maps using small sample numbers.

The presence of novel epigenetic marks in JIA neutrophils did not broadly reflect underlying genetic variation as determined from either whole genome sequencing or reference to previously identified JIA-associated SNPs/haplotypes. We could account for no more than 26% of the epigenetic variation, as reflected in histone marks, in JIA neutrophils by underlying genetic variation as reflected in indels (**S.Table7**). At the same time, we found that 31% of the LD blocks containing the disease-associated SNPs identified by Hinks et al [10] (see **Tables S1** and **S2**) also had an H3K4me1/H3K27ac-marked region seen only in neutrophils of either of children with untreated disease or in neutrophils of healthy children but not children with JIA, treated or untreated. The neutrophil epigenome may be independent of underlying genetic variation, although larger studies using combined genetics-genomics approches [38], are more likely to answer that question definitively. Our findings are consistent with recently published observations demonstrating that considerable variability in immune function in humans is determined by non-heritable factors[41]. Although there is clearly a measurable genetic component to JIA [10], and that genetic risk is located largely within the non-coding, functional genome [34], (a finding that JIA shares in common with other chronic inflammatory diseases [42]), our findings suggest the epigenetic differences we see between the neutrophils of children with JIA and those of healthy children may reflect environmental influences. These influences could include, of course, genetically mediated alterations in adaptive (T cell, B cell) immune function that are reflected in neutrophil epigenomes.

Our study pinpoints one environmental factor that is impacts neutrophil epigenomes: the initiation of effective therapy. This study is the first, to our knowledge, to demonstrate that effective therapy for a chronic human disease alters the epigenome. Furthermore, alterations in the epigenome associated with therapy tended to “correct” the neutrophil epigenome closer to patterns found in the cells of healthy children, as shown in Figure S3. Alterations in enhancer location included both poised (H3K4me1 marks) and active (H327ac marks) enhancers. Furthermore, even with this small number of samples, we were able to associate alterations in enhancer locations with alterations in gene expression. These findings support the idea that building genomic maps for each stage of therapy, from active disease through remission, as defined by the Wallace criteria [43] may be a useful approach to understanding the underlying biology of therapeutic response.

It was interesting to note that we did not see a strong correlation between changes in DNA methylation and changes in gene expression as they occurred with initiation of therapy in JIA neutrophils. There is good experimental evidence that DNA methylation does regulate gene expression in human neutrophils [44]. Our inability to identify a link between DNA methylation and gene expression with these samples very likely indicates an important limitation to this approach, which is likely to be insensitive to subtle links between epigenetic changes and gene expression. Under any circumstances, our experimental data suggest that the repositioning of enhancer marks has a stronger influence on gene expression than DNA methylation in the setting of therapeutic response in JIA, and support efforts to map important functional genomic elements in pathologically relevant cells in this disease.

Our data add another important element to the understanding of the genomic basis of JIA and other chronic inflammatory diseases. Multiple studies link genetic risk for chronic inflammatory diseases to previously annotated functional elements, including enhancers [34],[42]. The presence of “disease-specific enhancers” adds another layer of complexity to the idea that many chronic inflammatory diseases may essentially be disorders of transcriptional control. The coordination of transcription on a genome-wide basis is a complex process involving multiple elements that regulate chromatin accessibility, transcription factor binding, and transcription initiation [17]. Thus, sorting through the mechanisms through which genetic and epigenetic variation alter transcription needs to begin with the understanding that the epigenomes of peripheral blood cells (including the locations of functional elements) of patients with chronic inflammatory diseases are likely to be distorted compared to the reference ENCODE and/or Roadmap Epigenomics data.

### Conclusions

In this study, we demonstrate the feasibility of generating multidimensional genomic maps for a complex childhood disease from small numbers of samples, following the approaches used in the ENCODE and Roadmap Epigenomics projects. We demonstrate that such maps can be used to identify the presence of disease and disease-state associated epigenetic alterations in peripheral blood cells and the plasticity of the epigenome during therapy for JIA. We also demonstrate that transcriptional alterations linked to epigenetic alterations can be identified. We were unable to unambiguously identify genetic variants (SNPs, indels) that overlapped with alterations the JIA neutrophil epigenomes, possibly because this parsimonious method for mapping epigenomes is insensitive for detecting such overlap. Taken together, however, our findings demonstrate the feasibility of generating well-mapped epigenomes in pathologically relevant cells in disease states using the same approaches as were used in the Roadmap Epigenomics project. We expect that the generation of such maps in other cell types (CD4+ T cells, monocytes, NK cells, etc) will provide a wealth of new information re: the underlying pathobiology of complex inflammatory diseases like JIA.

## Acknowledgments of Funding

This work was supported by grants from the National Institutes of Health (R01-AI084200 and R01-AR060604 [JNJ), the Oklahoma Center for the Advancement of Science and Technology (HR07-139 [JNJ]), and an Innovative Research Grant #5989 from the Arthritis Foundation (JNJ). X.X, D.L., and T.W. are supported by NIH grants R01HG007354, R01HG007175, R01ES024992, U01CA200060, U24ES026699, and American Cancer Society Research Scholar grant RSG-14-049-01-DMC. This work was also supported by This work was also supported by the National Center for Advancing Translational Sciences of the National Institutes of Health under award number UL1TR001412 to the University at Buffalo. The funders had no role in study design, data collection and analysis, decision to publish, or preparation of the manuscript. The content is solely the responsibility of the authors and does not necessarily represent the official views of the NIH or the other funders.

## Author Contributions

Lisha Zhu – Was primarily responsible for the computational analysis and the writing of the manuscript.

Kaiyu Jiang – Performed the laboratory procedures for ChIPseq and RNAseq data and assisted in data analysis and interpretation.

Laiping Wong – Performed the computational analyses on the whole genome sequencing data and assisted in writing the manuscript.

Michael J. Buck – Provided guidance with the ChIP sequencing procedures.

Yanmin Chen - Assisted with the laboratory procedures for ChIPseq and RNAseq data.

Halima Moncrieffe, Laura McIntosh – Assisted in sample acquisition and preparation.

### Assisted in data interpretation

Tao Liu – Directed all aspects of the computational analysis

Xiaoyun Xing, Daofeng Li – Assisted with laboratory procedures and preliminary computational analysis of the methylation sequencing data.

Ting Wang – Directed the methylation sequencing.

James N. Jarvis – Designed the study. Assisted in data analysis and interpretation and writing of the manuscript.

---

**Figure S1.**
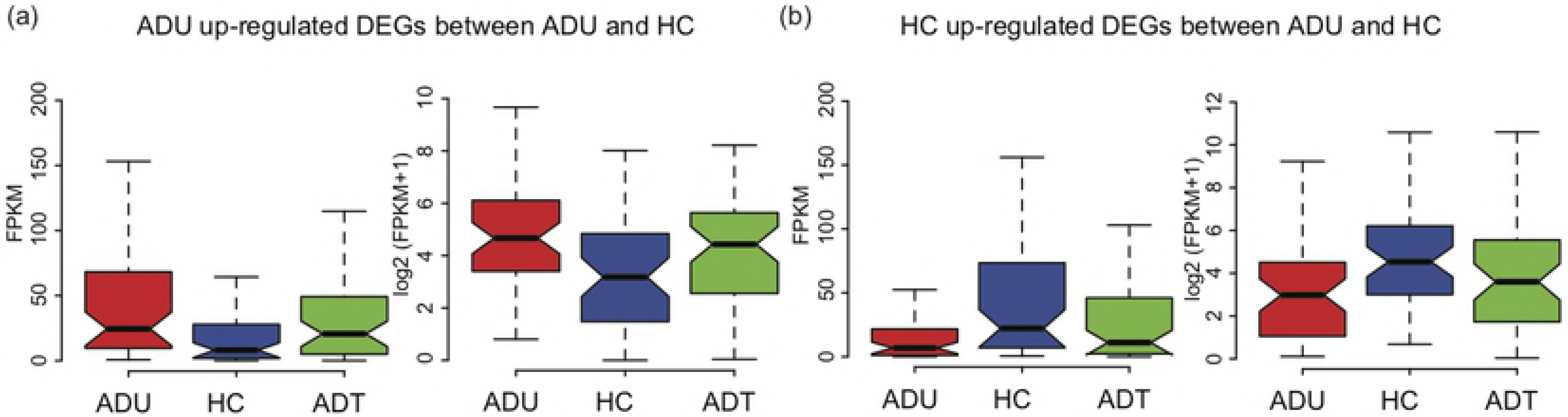
Gene expression levels of differentially expressed genes (DEGs) between ADU and HC in three groups. (a) boxplot of FPKM values of ADU up-regulated DEGs between ADU and HC in ADU, HC and ADT groups. (b) boxplot of FPKM values of ADU down-regulated DGEs between ADU and HC in ADU, HC and ADT groups.

**Figure S2.**
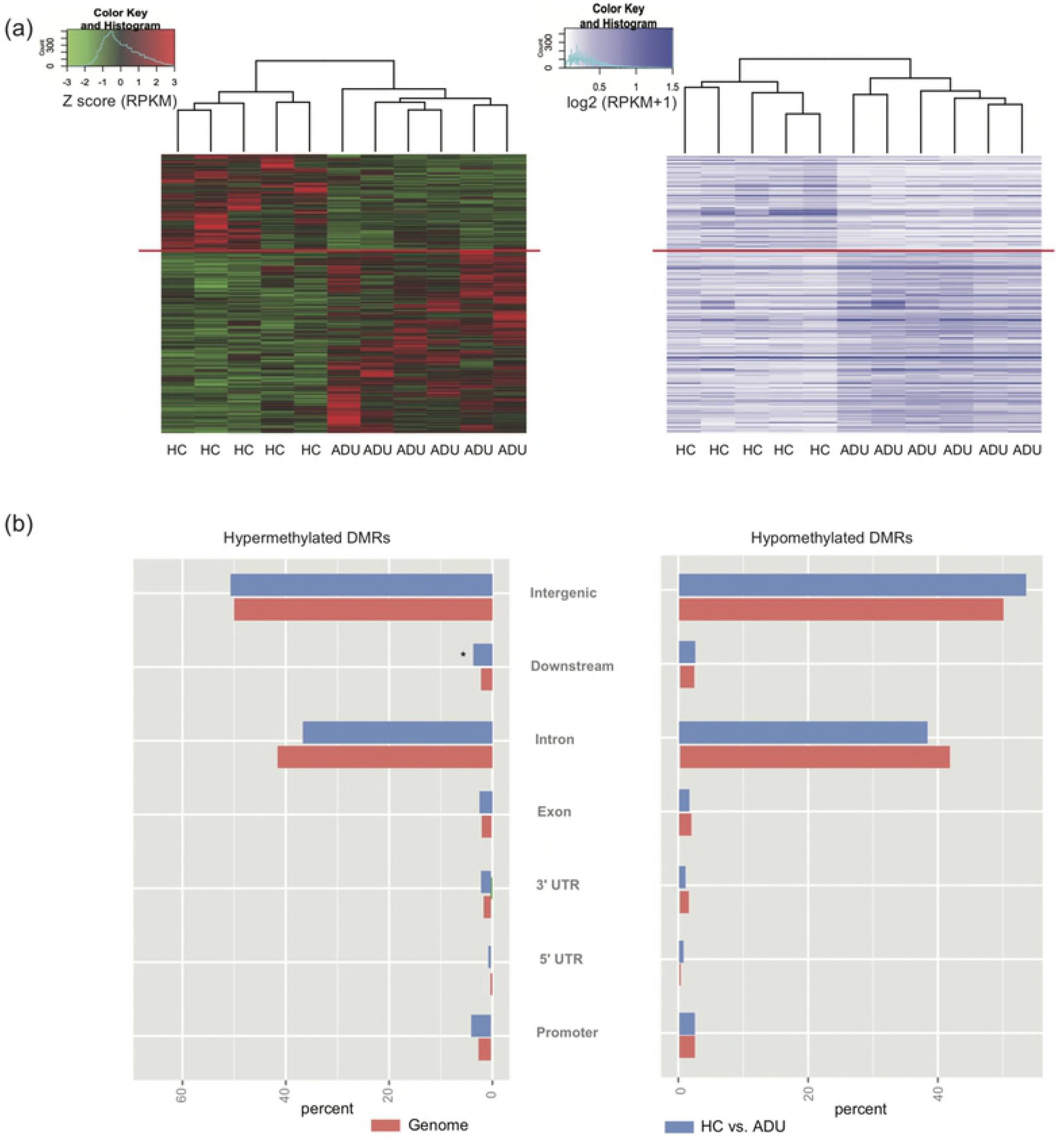
Heatmap and genomic distribution of DMRs between ADU and HC. (a) Heatmap of DMR RPKM values. Left: Z-score normalized values; Right: raw values. (c) Genomic distribution of hypermethylated and hypomethylated DMRs between ADU and HC. (Fisher’s exact test for the difference between DERs and genome background, * P < 10^−6^).

**Figure S3.**
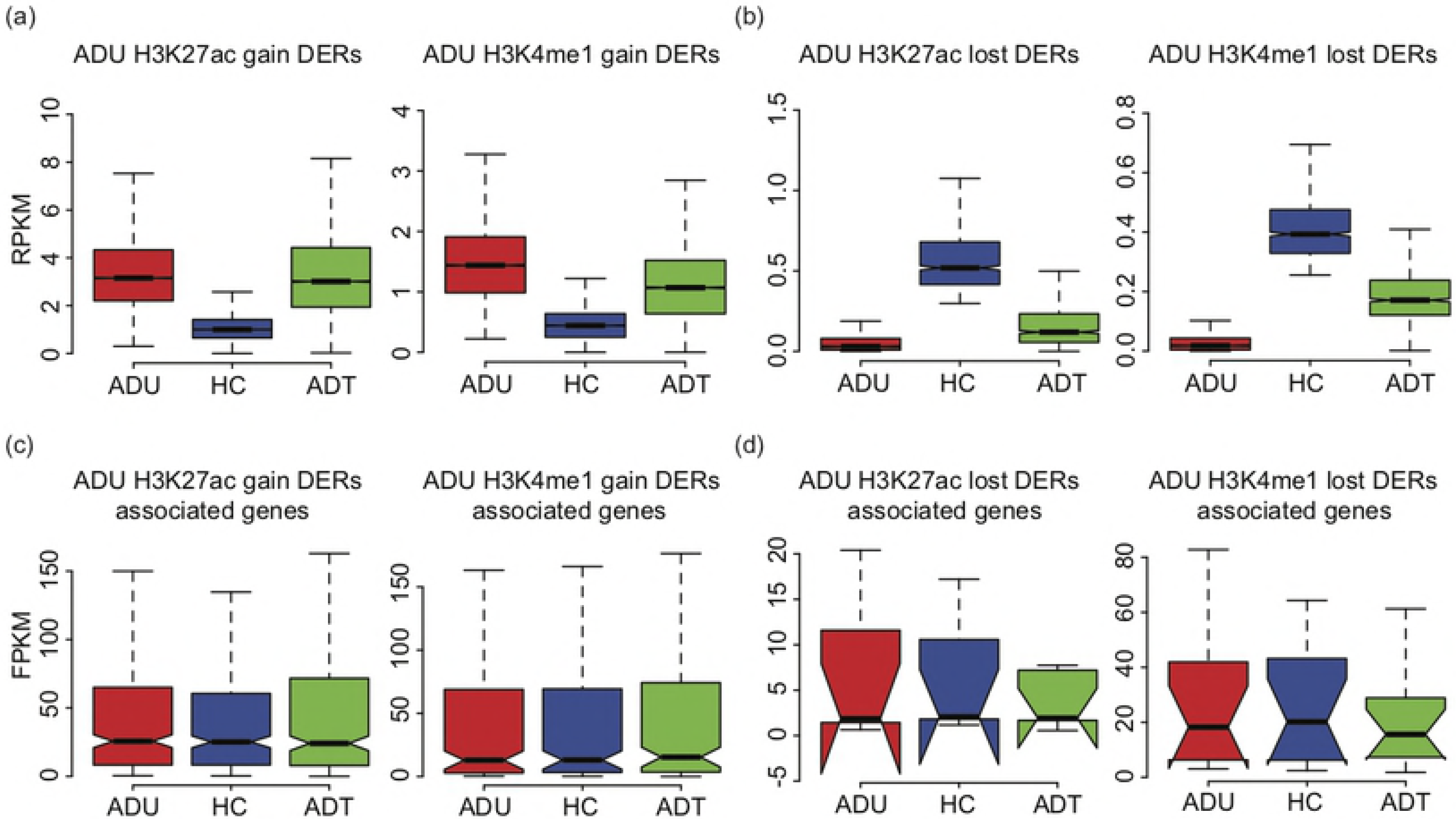
Treatment alters JIA neutrophil epigenomes. (a) boxplot of H3K27ac levels of total ADU H3K27ac gain DERs (left) and H3K4me1 levels total ADU H3K4me1 gain DERs (right) between ADU and HC in ADU, HC and ADT groups. (b) boxplot of H3K27ac levels of total ADU H3K27ac loss DERs (left) and H3K4me1 levels total ADU H3K4me1 loss DERs (right) between ADU and HC in ADU, HC and ADT groups. (c) boxplot of gene expressions in FPKM values of genes associated with ADU H3K27ac gain DERs (left) and ADU H3K4me1 gain DERs (right) that identified in the comparison between ADU and HC which also changed significantly between ADU and ADT. (d) boxplot of gene expressions in FPKM values of genes associated with ADU H3K27ac loss DERs (left) and ADU H3K4me1 loss DERs (right).

